# Tendon Cell Deletion of IKKβ/NF-κB Drives Functionally Deficient Tendon Healing and Altered Cell Survival Signaling In Vivo

**DOI:** 10.1101/2020.03.03.974774

**Authors:** Katherine T. Best, Emma Knapp, Constantinos Ketonis, Jennifer H. Jonason, Hani A. Awad, Alayna E. Loiselle

**Affiliations:** Center for Musculoskeletal Research, University of Rochester Medical Center, Rochester, NY 14642, United States of America; Department of Biomedical Engineering, University of Rochester, Rochester, New York, United States of America

**Author notes:** **Corresponding Author**, Alayna E. Loiselle, PhD, Center for Musculoskeletal Research, University of Rochester Medical Center, 601 Elmwood Ave, Box 665, Rochester, NY, 14642.

## Abstract

Acute tendon injuries are characterized by excessive matrix deposition that impedes regeneration and disrupts functional improvements. Inflammation is postulated to drive pathologic scar tissue formation, with nuclear factor kappa B (NF-κB) signaling emerging as a candidate pathway in this process. However, characterization of the spatial and temporal activation of canonical NF-κB signaling during tendon healing *in vivo*, including identification of the cell populations activating NF-κB, is currently unexplored. Therefore, we aimed to determine which cell populations activate canonical NF-κB signaling following flexor tendon repair with the goal of delineating cell-specific functions of NF-κB signaling during scar mediated tendon healing. Immunofluorescence revealed that both tendon cells and myofibroblasts exhibit prolonged activation of canonical NF-κB signaling into the remodeling phase of healing. Using cre-mediated knockout of the canonical NF-κB kinase (IKKβ), we discovered that suppression of canonical NF-κB signaling in Scleraxis-lineage cells increased myofibroblast content and scar tissue formation. Interestingly, Scleraxis-lineage specific knockout of IKKβ increased the incidence of apoptosis, suggesting that canonical NF-κB signaling may be mediating cell survival during tendon healing. These findings suggest indispensable roles for canonical NF-κB signaling during flexor tendon healing.

**One Sentence Summary:** Scleraxis-lineage specific knockdown of persistent canonical IKKβ/NF-κB drives scar formation and apoptotic signaling during flexor tendon healing.

## Introduction

Acute tendon injuries commonly heal with excessive scar tissue deposition, which complicates repair and drives functional deficits of the healing tendon. Scar-mediated tendon healing is especially problematic in flexor tendons of the hand as excessive scar tissue deposition can greatly reduce range of motion, negatively impacting patient quality of life(*1*). Both chronic inflammation and persistence of matrix-producing cells have been implicated as predominant drivers of scar-mediated healing(*2-6*); however, the cellular and molecular contributions during tendon healing are not well defined(*7*).

The canonical nuclear factor kappa B (NF-κB) signaling cascade drives both inflammatory and pro-survival gene expression, greatly affecting the outcomes of healing tissues(*8-11*). In short, the canonical heterodimer, most commonly a complex between proteins p50 and p65, is held in the cytoplasm by the inhibitor of κB (IκBα) protein, preventing translocation into the nucleus and subsequent transcriptional initiation of NF-κB-dependent genes. Through a wide variety of possible receptor-ligand interactions, the IκB kinase (IKK) complex, comprised of IKKα, IKKβ, and NEMO, becomes activated following phosphorylation of IKKβ, which subsequently phosphorylates and marks IκBα for degradation. IκBα degradation facilitates p50-p65 translocation into the nucleus, initiating NF-κB-dependent downstream gene expression.

Canonical NF-κB signaling has recently been implicated in scar-mediated tendon healing. Canonical NF-κB signaling activation is elevated in fibrotic human tendon tissue compared to healthy human tendon controls(*12*). In addition, global overactivation of canonical NF-κB signaling in a mouse model of acute flexor tendon injury and repair drives deposition of collagen matrix at the repair and increases the presence of alpha smooth muscle actin (αSMA+) myofibroblasts and F4/80+ macrophages, two cell types that jointly drive scar tissue formation(*2, 13*). Global inhibition of canonical NF-κB signaling via either p65-directed siRNA or a variety of inhibitors resulted in decreased scar/adhesion formation, diminished αSMA+ myofibroblast presence, and decreased collagen production(*12*). Additionally, cultured tendon fibroblasts treated with p65 inhibitors exhibited decreased proliferation and increased apoptosis, suggesting that canonical NF-κB signaling alters tendon cell survival(*12*). While informative, these studies do not examine the specific roles of NF-κB-activating cell populations during tendon healing *in vivo.* Recent work has demonstrated that deletion of IKKβ in tendon fibroblasts leads to decreased secretion of many pro-inflammatory factors, such as IL-6 and CCL2, compared to wildtype tendon cells in vitro(*14*). While Abraham *et al.* demonstrated that IKKβ deletion in *Scleraxis* (*Scx*)-lineage (Scx^Lin^) cells during enthesis healing resulted in improved biomechanical property outcomes relative to wildtypes(*14*), no study has assessed the cell-type-specific effects of canonical NF-κB signaling on tendon healing.

Therefore, in the present study we tested the hypothesis that flexor tendon cells activate canonical NF-κB signaling following an acute mid-substance tendon injury in vivo and these NF-κB-activated tendon cells contribute to excessive scar tissue formation during tendon healing. We characterized the spatial and temporal patterns of canonical NF-κB signaling in vivo and identified specific fibroblast populations (tendon cells, Scx^Lin^ cells, and myofibroblasts) that activate and maintain canonical NF-κB signaling following the inflammatory phase of healing, suggesting previously unappreciated roles for canonical NF-κB signaling in promoting the fibrotic response to tendon injury. Additionally, we assessed the effects of Scx^Lin^ specific IKKβdeletion (IKKβKO^Scx^) on tendon healing via gliding and biomechanics testing, analysis of signaling pathway activation, and histology. We also demonstrated that human tenolysis patient scar tissue had canonical NF-κB activated myofibroblasts present, suggesting that these findings are clinically relevant.

## Results

### Persistence of canonical NF-κB signaling during fibrotic tendon healing

To assess the spatial localization and temporal pattern of canonical NF-κB signaling throughout flexor tendon healing (days 3-28), C57Bl/6J FDL repair section were stained for phosphorylated p65 (p-p65), a marker of canonical NF-κB activation. A small number of peripheral scar tissue cells were p-p65+ at day 3 post-repair (Fig. 1A). The number of p-p65+ cells increased substantially by day 7 post-repair, primarily in cells residing within the developing scar tissue (Fig. 1A). Interestingly, p-p65+ cells persisted at the injury site through days 14 (Fig. 1A), 21 (Fig. 1A), and 28 post-repair (Fig. 1A), with a substantial portion of these cells residing within the native tendon stubs, suggesting that sustained activation of canonical NF-κB signaling in tendon fibroblast populations following the acute inflammatory phase of tendon healing. This prolonged activation of canonical NF-κB signaling, while primarily associated with acute inflammation, is consistent with recently published data suggesting involvement of inflammatory pathways well into the later phases of tendon healing(*15*).

**Figure 1.**
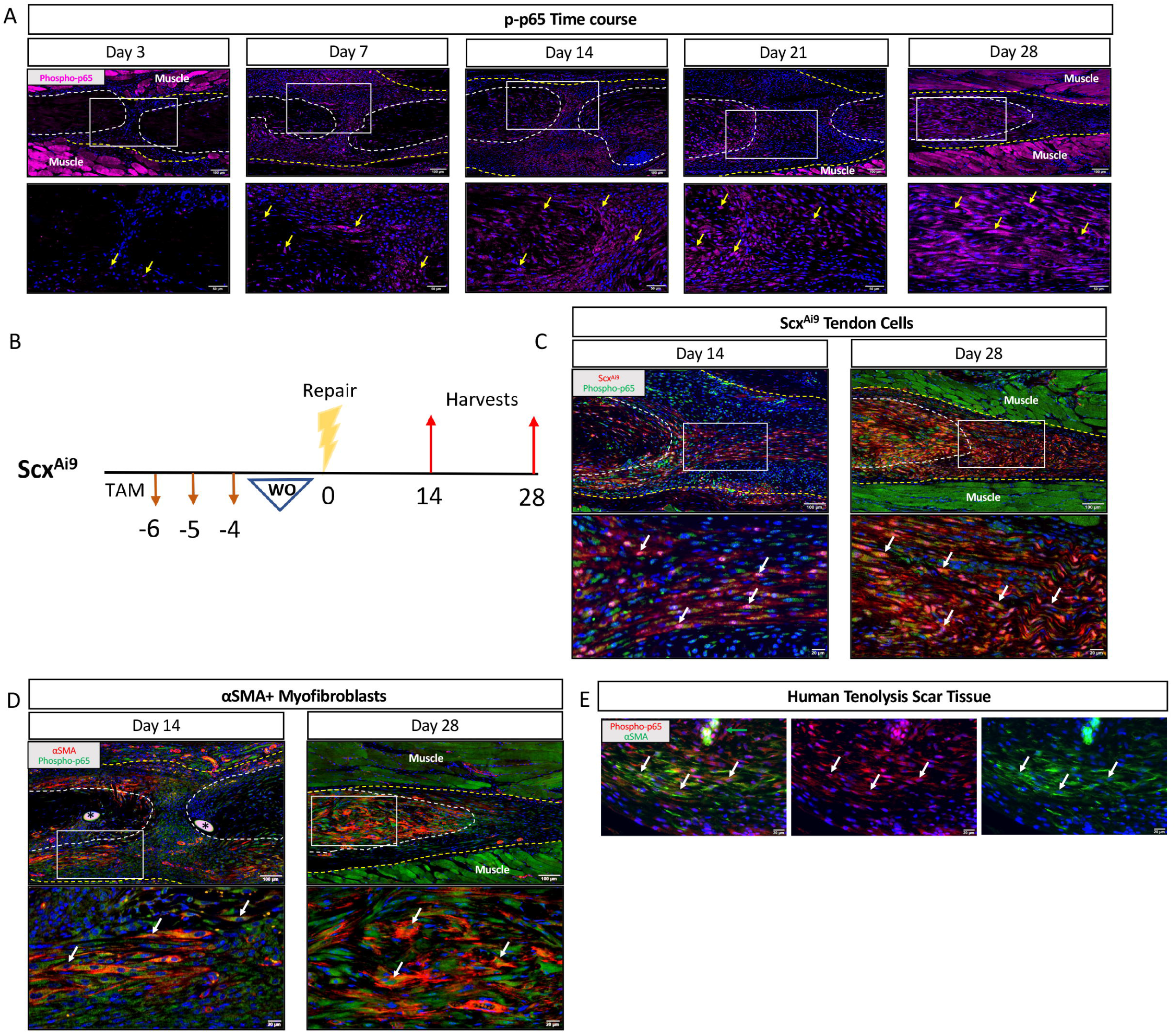
Canonical NF-κB signaling persists throughout tendon healing and is activated in tendon cells and αSMA+ myofibroblasts. Phosphorylated p65 (p-p65) is a marker of canonical NF-κB signaling. Immunofluorecence of p-p65 in C57Bl/6J mice at 3, 7, 14, 21, and 28 days post-repair (A). Examples of positive stain indicated by yellow arrows. Scx-Cre^ERT2^ mice were crossed to the ROSA-Ai9 reporter to label adult Scx tendon cells throughout healing (Scx^Ai9^) (B). Co-immunofluorescence of either tdTomato (labels Scx^Ai9^) (C) or αSMA (D) with p-p65 in Scx^Ai9^ or C57Bl/6J mice, respectively. Tendon is outlined by white dotted line and scar tissue by yellow dotted line. White boxes indicate location of higher magnification images. Examples of co-localization indicated by white arrows. Sutures indicated by *. N=3-4. Coimmunofluorescence of αSMA and p-p65 in human tenolysis scar tissue (E). Co-localization indicated by white arrows and blood vessel by green arrow. Nuclei stained blue with DAPI. Auto-fluorescent muscle is labeled with “Muscle”.

### Adult tendon cells and myofibroblasts activate and maintain canonical NF-κB signaling

The presence of p-p65+ cells within the tendon stubs following repair suggested that tendon fibroblast populations may activate canonical NF-κB signaling during healing. To assess canonical NF-κB activation specifically in tendon cells, Scx^Ai9^ mice were injected with tamoxifen (TAM) for three days, with four days between the final injection and the day of surgery to ensure washout of TAM (Fig. 1B). Co-immunofluorescence between tdTomato (labels Scx^Ai9^ cells) and p-p65 at days 14 and 28 post-repair confirmed that tendon cells are a predominant population activating canonical NF-κB signaling during post-inflammatory phases of healing (Fig. 1C). We have previously shown that myofibroblasts, a contractile cell type that can contribute to both proper healing and pathologic tissue fibrosis(*3, 16*), are present at the healing tendon(*17, 18*). Co-immunofluorescence between the myofibroblast marker αSMA and p-p65 demonstrated that myofibroblasts activated canonical NF-κB signaling at both day 14 and 28 post-repair (Fig. 1D). Intriguingly, clinical tendon scar tissue obtained during tenolysis surgery also contained p-p65-expressing αSMA+ myofibroblasts, demonstrating the clinical relevance of sustained NF-κB activation in the fibrotic response to tendon injury (Fig. 1E).

### Scleraxis-lineage cells label both tendon cells and myofibroblasts during tendon healing

We have previously shown that most *Scx*-expressing tendon cells do not differentiate into αSMA+ myofibroblasts(*17*). However, the myofibroblast fate of Scx^Lin^ cells has not been assessed. While Scx^Ai9^ mice label all cells actively expressing *Scx* at the time of TAM injection (nearly 70% of tendon cells(*17*)), Scx^Lin^ mice label all cells derived from the *Scx* lineage. The majority of tendon cells are Scx^Lin+^in uninjured mouse flexor tendon (Fig. 2A). Scx^Lin+^ cells are found throughout the scar tissue at day 7 post-repair (Fig. 2A). At days 14 and 21, Scx^Lin+^ cells have organized into a cellular bridge spanning the scar tissue between the tendon stubs (Fig. 2A), similar to what was seen with Scx^Ai9^ mice using the flexor tendon repair model(*17*). Virtually all αSMA+ myofibroblasts present at the healing tendon are Scx^Lin+^ and p-p65+ and days 14 (Fig. 2B) and 28 (Fig. 2C) post-repair. Previous work has suggested that these are recruited αSMA+ cells that subsequently activate Scx expression post-repair(*19*). Therefore, Scx-Cre mice provide an excellent opportunity to modulate canonical NF-κB signaling in both the tendon cell and myofibroblast populations during tendon healing to understand how canonical NF-κB signaling in fibroblast populations modulate healing.

**Figure 2.**
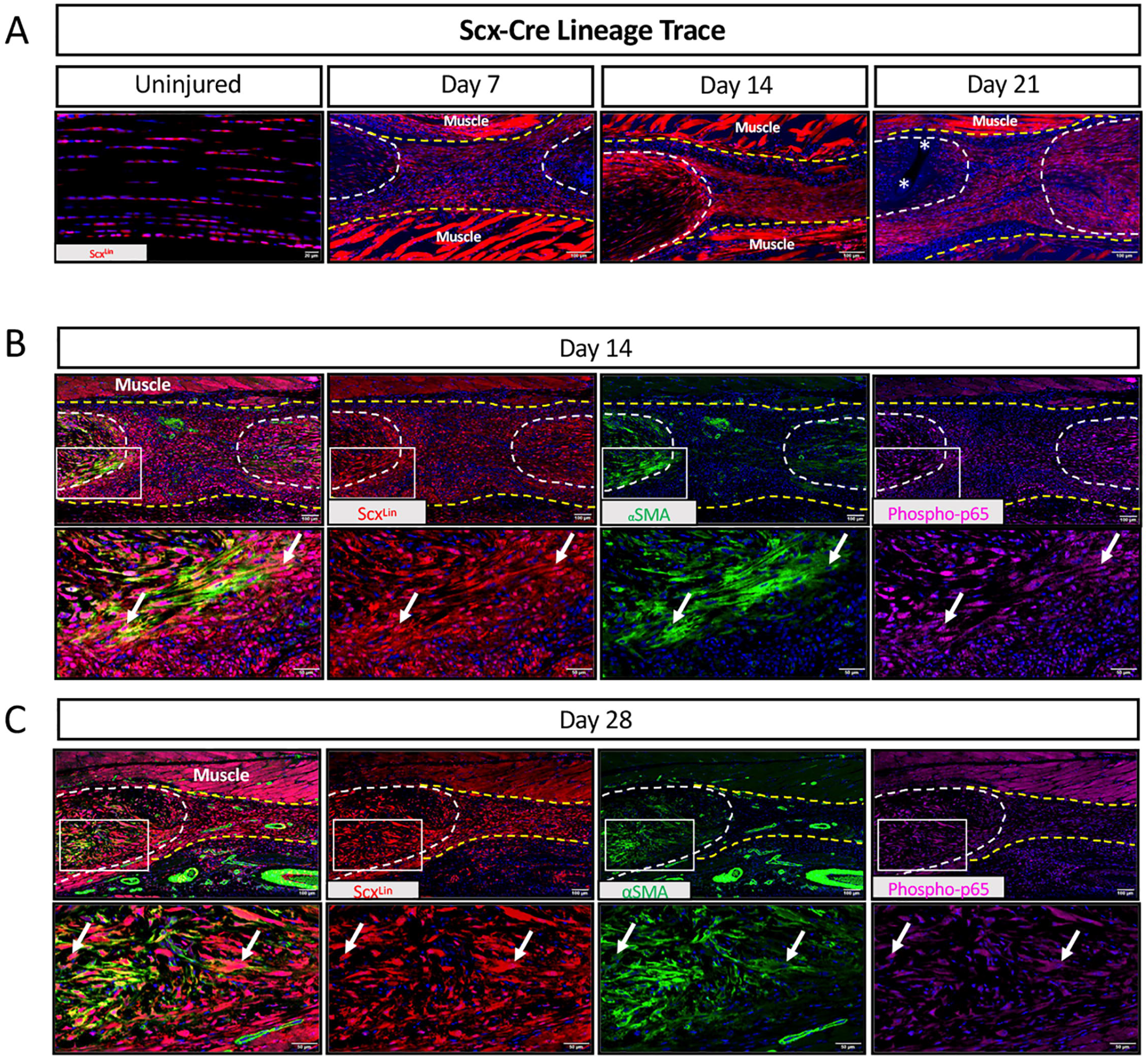
Scleraxis-Cre mice can be used to target both tendon cells and myofibroblasts. Scx-Cre mice were crossed to the ROSA-Ai9 reporter to trace Scx lineage cells at homeostasis and throughout healing (Scx^Lin^). Hindpaws were harvested uninjured, and at days 7, 14, and 21 post-repair (A). Co-immunofluorescence between tdTomato (labels Scx^Lin^), αSMA, and p-p65 in Scx^Lin^ mice at days 14 (B) and 28 (C). Tendon is outlined by white dotted line and scar tissue by yellow dotted line. White boxes indicate location of higher magnification images. Examples of positive stain indicated by white arrows. Nuclei stained blue with DAPI. Auto-fluorescent muscle is labeled with “Muscle”. Sutures labeled by *. N=3-4.

### IKKβKO^Scx^ does not impact baseline tendon function or biomechanical properties

IKKβ is the predominant kinase driving downstream canonical NF-κB signaling (Fig. 3A). Scx-Cre animals were crossed to IKKβ^F/F^ mice to generate animals lacking either one (IKKβHet^Scx^) or both (IKKβKO^Scx^) copies of the NF-κB kinase IKKβ gene *IKK2* in Scx^Lin^ cells. Uninjured FDL tendons from IKKβHet^Scx^ and IKKβKO^Scx^ mice exhibited decreased levels of IKKβ protein compared to WT littermates (Fig. 3B). To confirm that IKKβKO^Scx^ did not significantly alter baseline tendon function or mechanical properties compared to WT, uninjured tendons were assessed. There were no significant differences detected between MTP flexion angle (Fig. 3C), gliding resistance (Fig. 3D), stiffness (Fig. 3E), or maximum load at failure (Fig. 3F) between genotypes.

**Figure 3.**
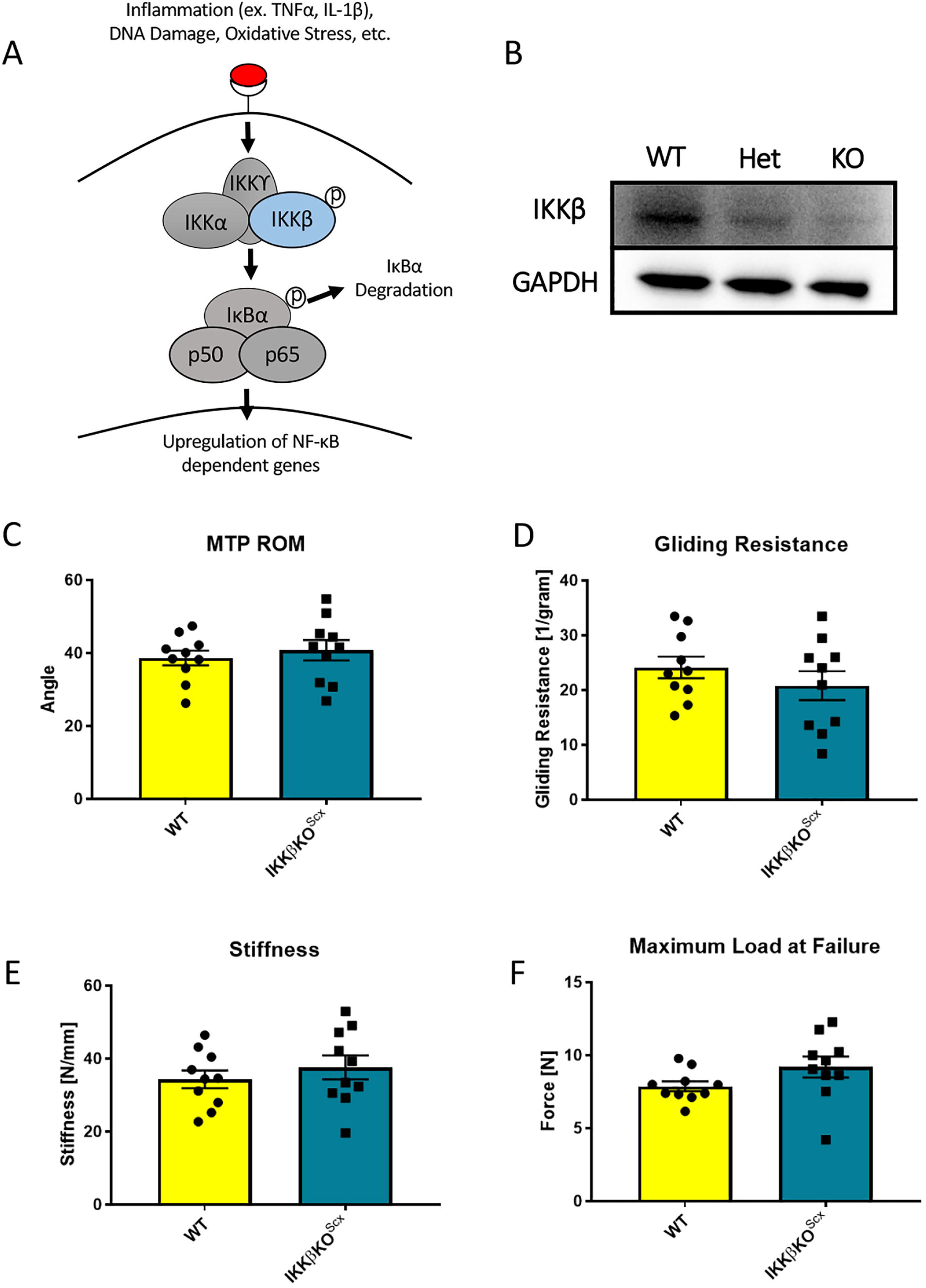
IKKβKO^Scx^ does not negatively affect baseline tendon gliding function or biomechanical propertes. IKKβ is the canonical kinase of NF-κB signaling that drives downstream gene expression (A). Western blot assessing IKKβ using protein extracted from WT, IKKβHet^Scx^ and IKKβKO^Scx^ uninjured tendons (B). N=3 tendons per genotype. Full length blot available in supplementary figure 1. Quantification of band intensity relative to GAPDH load control presented in supplementary table 1. Measurement of metatarsophalangeal (MTP) joint flexion angle (C), gliding resistance (D), stiffness (E), and maximum load at failure (F) of uninjured WT and IKKβKO^Scx^ tendons. N=10 per genotype. Students t-test used to assess statistical significance between genotypes.

### IKKβKO^Scx^ negatively impacts repair tendon gliding function by day 28 post-repair

MTP range of motion was significantly decreased in IKKβKO^Scx^ healing tendons relative to WT littermates at 28 days post-repair (WT: 36.44 ± 4.04, KO: 25.61 ± 2.95, p = 0.0368) (Fig. 4A). Between days 14 and 28, WT tendon MTP range of motion improved 41.35% while IKKβKO^Scx^ tendons only improved 3.31%. Gliding resistance was significantly increased in IKKβKO^Scx^ tendons relative to WT littermates at day 28 post-repair (WT: 26.34 ± 4.89, KO: 47.36 ± 8.80, p = 0.0384) (Fig. 4B). Between days 14 and 28, WT tendon gliding resistance decreased 33.25% while IKKβKO^Scx^ tendons increased by 7.20%. IKKβKO^Scx^ tendon repairs did not exhibit significantly different stiffness or maximum load at failure at either day 14 or 28 post-repair relative to WT littermates (Fig. 4C,D).

**Figure 4.**
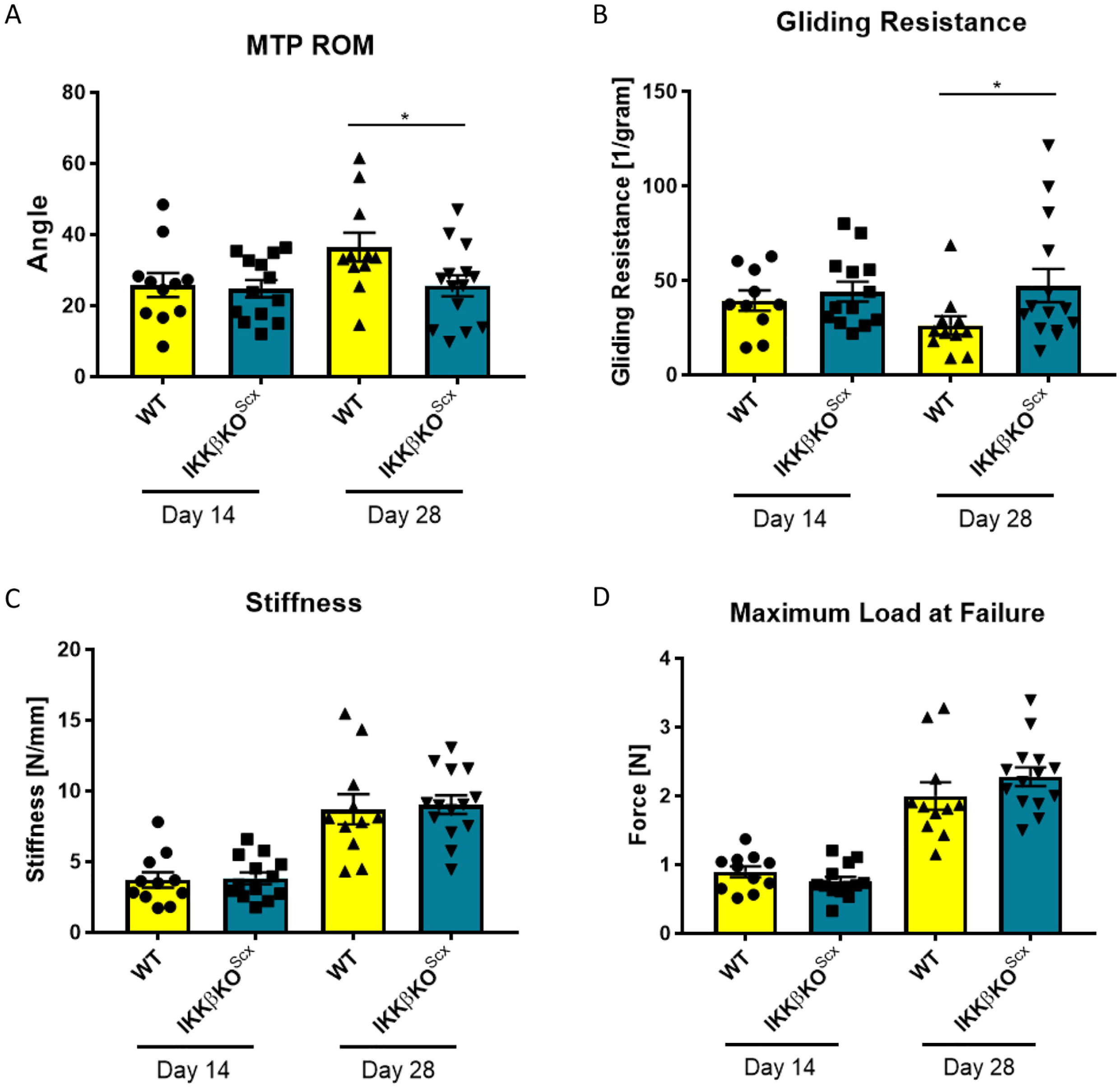
IKKβKO^Scx^ heal with impaired gliding ability. Measurement of metatarsophalangeal (MTP) joint flexion angle (A), gliding resistance (B), stiffness (C), and maximum load at failure (D) of 14- and 28-day post-repair WT and IKKβKO^Scx^ tendons. N=10-14 per genotype. Students t-test was used to assess statistical significance between genotypes at a given time point, except for gliding resistance which required a Mann-Whitney test. * indicates p < 0.05

### IKKβKO^Scx^ heal with increased matrix deposition

As IKKβKO^Scx^ repairs exhibited trending deficits in gliding ability, we next wanted to assess if tissue morphology or matrix deposition was affected. No differences in overall tissue morphology were detected using Alcian blue hematoxylin orange G stain (Fig. 5A). While there were no apparent differences in collagen content at day 14, IKKβKO^Scx^ repairs exhibited regions of disorganized collagen matrix at the tendon-muscle border that was not detected in WT littermates at day 28 (Fig. 5B, black arrows). In addition to collagen, the pro-fibrotic matricellular protein periostin was assessed. While no differences in periostin were detected at day 14 (Fig. C, C’), IKKβKO^Scx^ repairs had significantly increased periostin at day 28 post-repair (D14: WT vs. KO, p = 0.4353; D28: WT vs. KO, p = 0.0084) (Fig. C, C”)(*20*).

**Figure 5.**
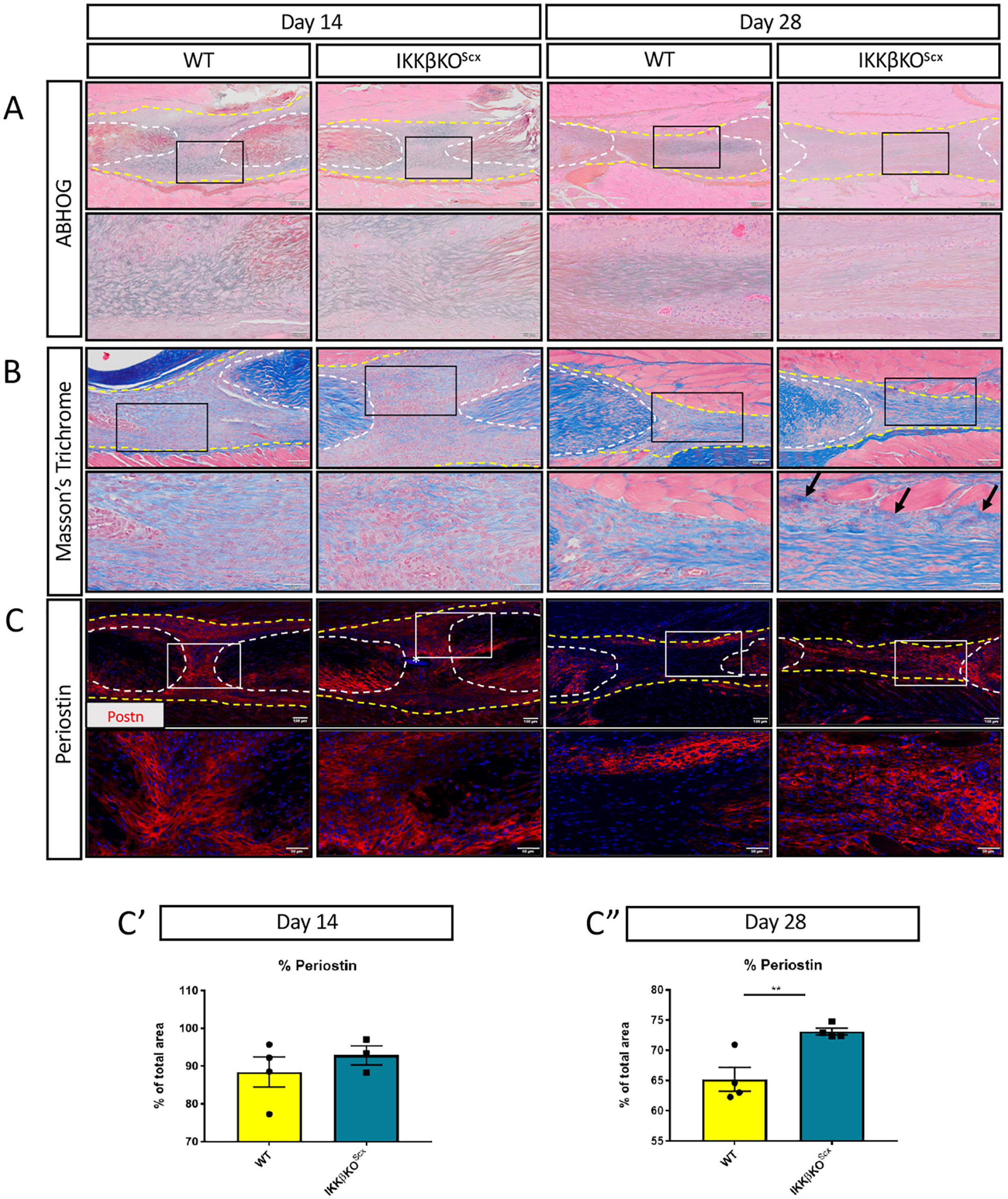
IKKβKO^Scx^ repairs heal with disorganized collagen and increased periostin. Histology of WT and IKKβKO^Scx^ tendons 14 and 28 days post-repair. Alcian blue/hematoxylin and Orange G stain utilized to assess overall morphology (A). Masson’s trichrome stain used to visualize collagen content and organization (B). Example of collagen disorganization indicated by black arrow. Tendon is outlined by white dotted lines and scar tissue by yellow dotted lines in histology images. Immunofluorescence of periostin in WT and IKKβKO^Scx^ tendons 14 and 28 days post-repair (C). Tendon is outlined by white dotted line and scar tissue by yellow dotted line in immunofluorescent images. Nuclei stained blue with DAPI. Black (A,B) or White (C) boxes indicate location of higher magnification images. Sutures labeled by *. Quantification of periostin at 14 (C’) and 28 (C”) days post-repair. N=3-4. Student’s t-test used to assess statistical significance between genotypes at a given time point. ** indicates p < 0.01.

### IKKβKO^Scx^ alters S100a4+ cell and αSMA+ myofibroblast presence by day 14

We have previously described how global activation of canonical NF-κB signaling drives increased presence of macrophages and myofibroblasts at the site of healing tendon(*13*). Therefore, we wanted to assess if IKKβKO^Scx^ resulted in altered presence of these cell types during tendon healing. There was a trend towards increased F4/80+ macrophages in IKKβKO^Scx^relative to WT at 14 and 28 days post-repair (D14: WT vs KO, p = 0.0557, D28: WT vs KO, p = 0.2000), but these differences were not significant (Fig. 6A,D). IKKβKO^Scx^ healing tendons exhibited a significant, transient increase in αSMA+ myofibroblasts 14 days post-repair relative to WT repairs (D14: WT vs KO, p = 0.0338) that was no longer seen by day 28 (D28: WT vs KO, p = 0.3586) (Fig. 6B,D). Notably, between days 14 and 28 the presence of αSMA+ cells decreased by 5.10% in WT repairs and 28.44% in IKKβKO^Scx^ repairs, highlighting a rapid decline in overall myofibroblast content in IKKβKO^Scx^ healing tendons. We have previously established that S100a4 is a key driver of fibrotic tendon healing(*18*). S100a4+cell presence was also significantly increased at day 14 post-repair in IKKβKO^Scx^ healing tendons relative to WT littermates (D14: WT vs KO, p = 0.0240), while this difference was no longer seen by day 28 (D28: WT vs KO, p = 0.4641) (Fig. 6C,D).

**Figure 6.**
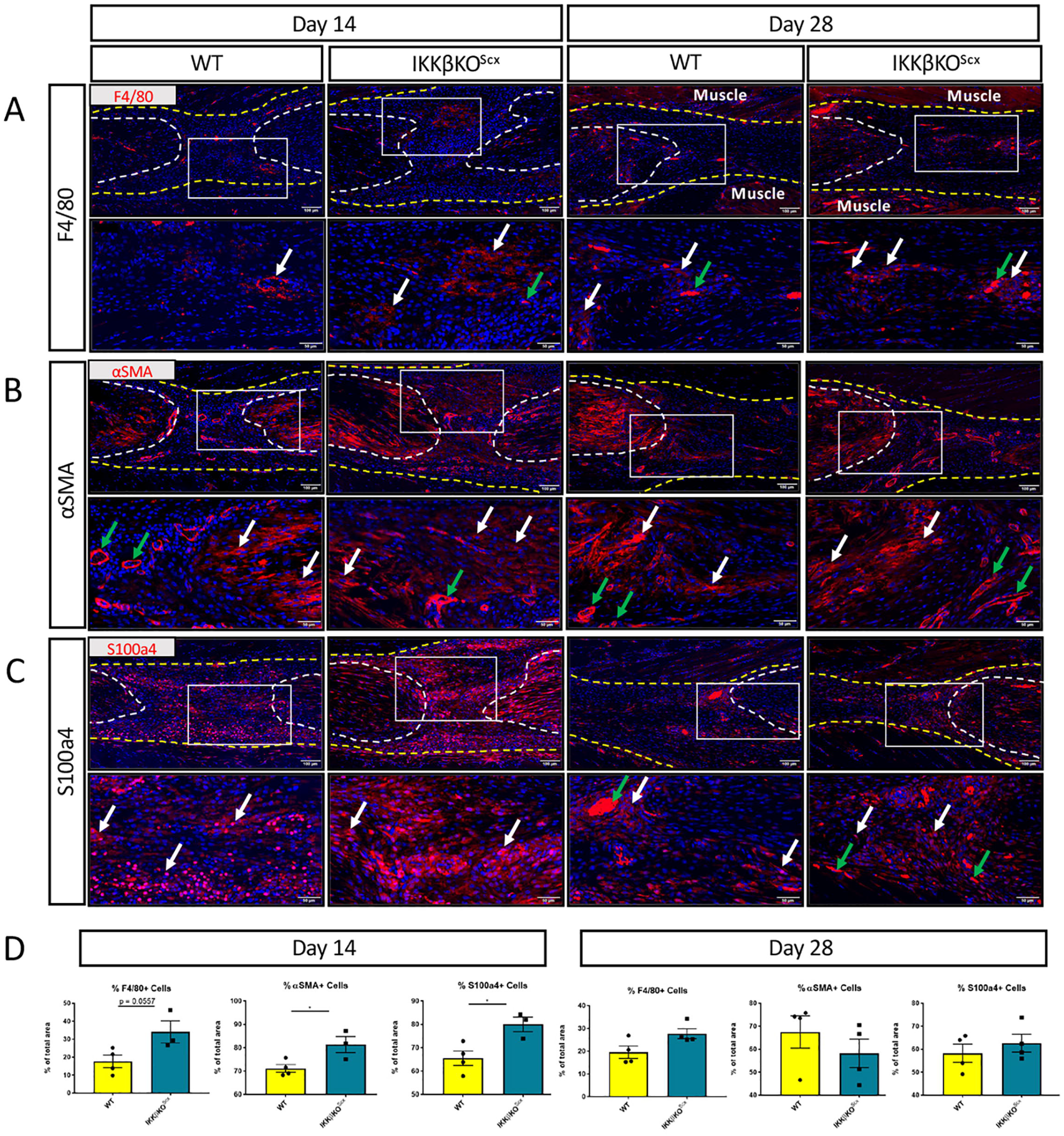
IKKβKO^Scx^ repairs heal with an altered cellular landscape compared to WT littermates. Immunofluorescence of WT and IKKβKO^Scx^ tendons 14 and 28-days post-repair to assess F4/80+ macrophages (A), αSMA+ myofibroblasts (B), and S100a4+ cells (C). Tendon is outlined by white dotted line and scar tissue by yellow dotted line. White boxes indicate location of higher magnification images. Where applicable, examples of positive stain indicated by white arrows, while examples of auto-fluorescent blood cells and α-SMA+ blood vessels indicated by green arrows. Auto-fluorescent muscle is labeled with “Muscle”. Quantification of F4/80, αSMA, and S100a4 fluorescence at days 14 and 28 post repair (D). N=3-4. Student’s t-test used to assess statistical significance between genotypes at a given time point, except for D28 F4/80 and αSMA which required a Mann-Whitney test. * indicates p < 0.05.

### IKKβKO^Scx^ modulates various signaling cascades during tendon healing

IKKβ is involved in a variety of signaling cascades(*21*). To assess how IKKβKO^Scx^ affects pathway activation relative to WT littermates at days 14 and 28 post-repair, western blotting was utilized. IKKβKO^Scx^ healing tendons exhibited decreased canonical NF-κB signaling (p-p65) relative to WT at day 14, as expected (Fig. 7A). Interestingly IKKβKO^Scx^ healing tendons also contained less total p65 relative to WT littermates (Fig. 7A). While ERK signaling was largely unaffected (Fig. 7A), p38 signaling cascades exhibited a transient increase in signaling at day 14 that was subsequently decreased by day 28 compared to wildtype controls (Fig. 7B). JNK signaling was suppressed at day 14 in IKKβKO^Scx^ repairs but were unchanged between genotypes at day 28 (Fig. 7B, Supplemental Table 2). Interestingly, IKKβKO^Scx^ healing tendons had elevated levels of total JNK at day 14 relative to wildtype littermates and did not undergo the increase in total JNK between days 14 and 28 exhibited by their wildtype littermates (7B). Both AKT and β-Catenin signaling pathways were decreased in IKKβKO^Scx^ healing tendons at 14 and 28 days post-repair, while S6K1 (mTOR signaling) was decreased in IKKβKO^Scx^ at day 28 compared to wildtypes (Fig. 7C,D). To assess the effect of IKKβKO^Scx^ on apoptosis, phospho-Foxo3a was examined. Decreased levels of phospho-Foxo3a were present in IKKβKO^Scx^ tendon repairs at both 14 and 28 days post-repair, indicative of increased apoptosis (Fig. 7D).

**Figure 7.**
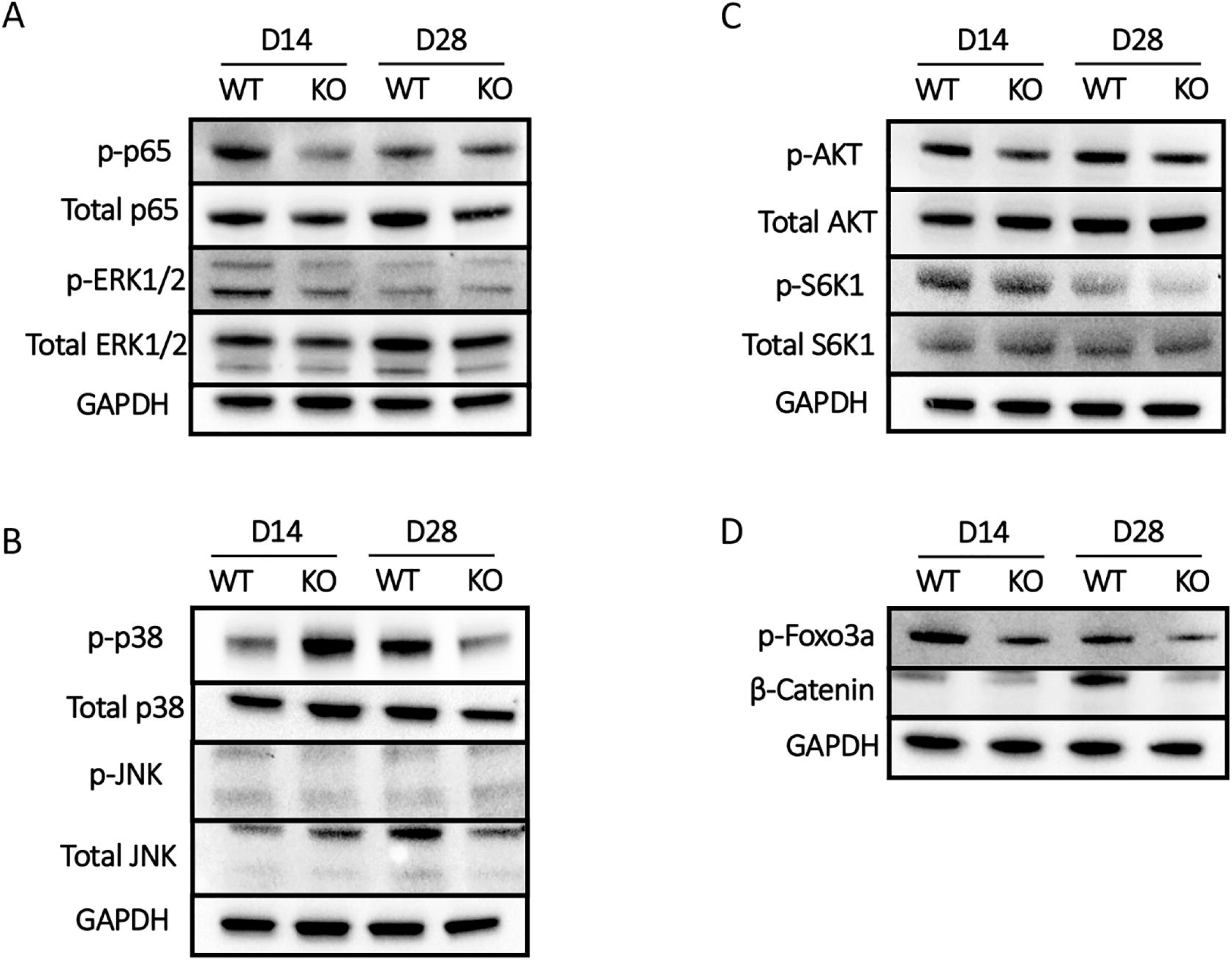
IKKβKO^Scx^ modulates several signaling cascades beyond canonical NF-κB signaling. Western blots to assess canonical NF-κB [p65, (A)], MAPK [ERK1/2 (A), JNK (B), p38 (B)], mTOR [S6K1 (C)], AKT (C), β-Catenin (D), and apoptosis [p-Foxo3a (D)] at 14 and 28 days post-repair in WT and IKKβKO^Scx^ tendons. Full length blots are presented in supplementary figure 2. Quantification of band intensity relative to GAPDH load control presented in supplementary table 2. N=3 tendons per genotype per timepoint.

### IKKβKO^Scx^ alters cell survival and drives increased apoptosis

Given the decreased p-Foxo3a observed in KKβKO^Scx^ tendon repairs, we further investigated the effects of IKKβ-dependent signaling on cell survival during healing, via assessment of NF-κB-associated survival factors Bcl-2 and Bcl-xL in wildtype and IKKβKO^Scx^ healing tendons 28 days post-repair. While not significant, there was a trending decrease in Bcl-2+ cells in IKKβKO^Scx^ repairs relative to wildtype littermates (WT vs KO, p = 0.0654) (Fig. 8A). No difference in Bcl-xL was detected between genotypes (Fig. 8B). Cleaved caspase 3 was also assessed as a marker for apoptotic cells. There was a significant increase in cleaved caspase 3+ apoptotic cells in IKKβKO^Scx^ repairs relative to wildtypes (WT vs KO, p = 0.0311) (Fig. 8C). Together this suggests that IKKβ signaling in Scx^Lin^ mediates cell survival, possibly through NF-κB dependent Bcl-2 expression.

**Figure 8.**
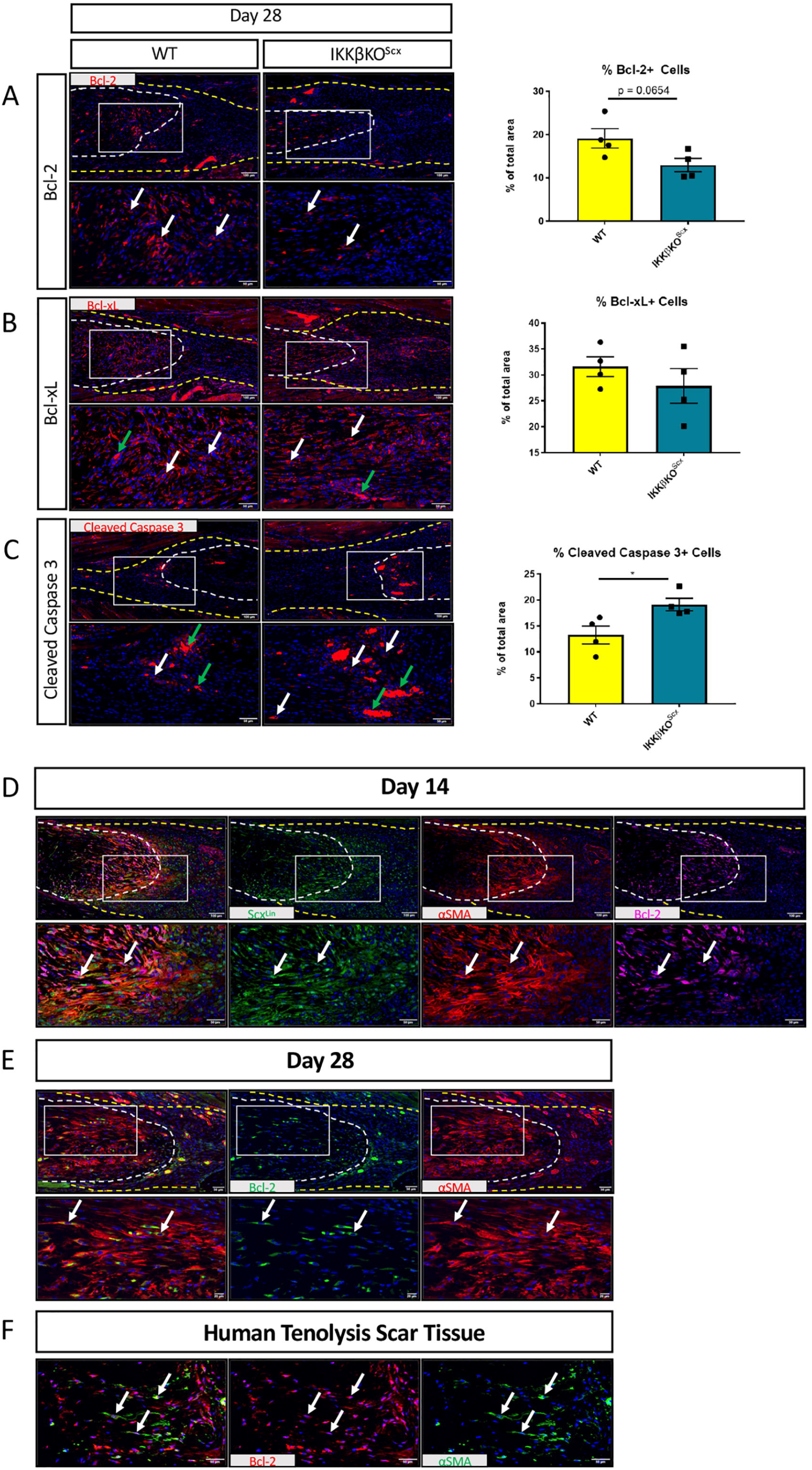
IKKβKO^Scx^ drives increased apoptosis. Immunofluorescence and quantification of WT and IKKβKO^Scx^ tendons 28-days post-repair to assess Bcl-2+ (A), Bcl-xL+ (B), and Cleaved Caspase 3+ cells (C). Tendon is outlined by white dotted line and scar tissue by yellow dotted line. White boxes indicate location of higher magnification images. Examples of positive stain indicated by white arrows, while examples of auto-fluorescent blood cells indicated by green arrows. N=4. Co-immunofluorescence between tdTomato (Scx^Lin^), αSMA, and BCL2 at day 14 post-repair mouse tendon (D), αSMA and BCL2 at day 28 post-repair in C57Bl/6J mouse tendon (E), and αSMA and Bcl-2 in human Tenolysis scar tissue (F). N=3-4 for mouse data. Student’s t-test used to assess statistical significance between genotypes at a given time point. * indicates p < 0.05.

We then confirmed that Bcl-2 is expressed by Scx^Lin^αSMA myofibroblasts at 14 days post-repair in the healing tendon (Fig. 8D). Additionally, Bcl-2 expression is maintained by αSMA myofibroblasts through day 28 post-repair (Fig. 8E). These results are in concordance with observations in human Tenolysis scar tissue in which αSMA myofibroblasts also express BCL-2 indicating the clinical relevance of these findings (Fig. 8F).

## Discussion

In the present study we characterized both the spatial localization and cell-type specific temporal activation profile of canonical NF-κB signaling following acute flexor tendon injury and repair. We determined that canonical NF-κB activation persists beyond the acute inflammatory phase of healing and is specifically activated by both tendon cells and myofibroblast populations during later phases of healing. Contrary to our hypothesis, IKKβ knockout in Scx^Lin^ cells resulted in increased scar formation which was accompanied by significant increases in S100a4+ cells and αSMA+ myofibroblasts detected during repair, disorganized collagen, increased periostin, and decreased cell survival (increased apoptosis).

It was recently demonstrated that IKKβ deletion in Scx^Lin^ cells resulted in improved biomechanical properties outcomes of the enthesis relative to wildtypes(*14*). The present study demonstrates that inhibition of canonical NF-κB signaling in Scx^Lin^ fibroblast populations *in vivo* is not biomechanically beneficial to flexor tendon healing and drives a pro-fibrotic response, suggesting that canonical NF-κB signaling may impact enthesis and flexor tendon healing in different ways. It has previously been shown that global overactivation of canonical NF-κB signaling drives increased matrix formation(*13*) while global inhibition of canonical NF-κB decreases matrix deposition and adhesion formation(*12*). There are several possibilities that could explain this discrepancy. First, IKKβKO^Scx^ exhibit decreased canonical NF-κB signaling via p-p65, suggesting decreased overall inflammation during healing. This likely results in a quicker resolution of the inflammatory phase of healing resulting in an earlier transition into the more proliferative and remodeling-focused stages of healing. Earlier proliferation of fibroblasts and differentiation of myofibroblasts could account for the increased presence of αSMA+ myofibroblasts at day 14 in IKKβKO^Scx^ healing tendon relative to WT and may drive excess scar formation during healing. IKKβKO^Scx^ cells in the healing tendon may also influence proliferation and recruitment patterns of both tendon cells and extrinsically recruited cells, resulting in a fundamentally altered cellular landscape at the repaired tendon compared to WT littermates, resulting in a more pro-fibrotic population of cells present at the repair. While changes in proliferation were not assessed in this study, there are clear changes in the cellular composition of the repair site between WT and IKKβKO^Scx^. Additionally, the loss of IKKβ in Scx^Lin^ cells may intrinsically alter functions of the Scx^Lin^ cells themselves, resulting in increased proliferation or prevalence of myofibroblast differentiation.

Inflammation has been implicated as the critical driver of excessive scar tissue formation, or fibrosis, in a variety of tissues(*22*). Inflammation is characterized by an influx of inflammatory cells such as neutrophils and macrophages. Macrophages have a variety of functions during healing. Notably, they secrete cytokines and chemokines that recruit and activate fibroblast cell populations, the cells primarily responsible for laying down matrix necessary for repair(*4*). Macrophages are also involved in driving fibroblast-myofibroblast differentiation(*2*). Myofibroblasts have historically been viewed as a specialized contractile fibroblast cell that can lay down matrix and aiding in wound closure. Chronic persistence of either macrophages or myofibroblast cells can result in continued, excessive deposition of matrix proteins, resulting in a fibrotic healing tissue(*2, 3, 5*). While canonical NF-κB signaling is often viewed as an immunecell activated inflammatory signaling cascade, it has previously been shown that fibroblast populations can activate NF-κB signaling to contribute to inflammation and pro-survival of myofibroblasts(*10, 14*), therefore promoting fibrotic healing. In this study we have shown that tendon and myofibroblast cell populations activate canonical NF-κB signaling during post-inflammatory phases of healing. Additionally, we have provided evidence that IKKβ-mediated signaling can influence cell survival during tendon healing *in vivo*, with IKKβKO^Scx^ resulting in increased apoptosis. Future studies investigating the influence of canonical NF-κB signaling on tendon cell and myofibroblast survival as a mechanism of fibrotic healing is an important future direction to explore, especially considering our data demonstrating that p-p65+ and Bcl-2+ myofibroblasts are present in human tendon scar tissue (Figs. 1 and 8).

While no changes in baseline tendon function were observed between genotypes, some IKKβKO^Scx^ mice presented with developmental abnormalities. A small number of male and female IK KβKO^Scx^ mice exhibited skin with rough scabby patches that became increasingly lesion-proned as they aged. While it has previously been shown that *Scx* is not detected in the skin(*23*), *Scx* is expressed in hair follicle cells(*24*) and keloid scars of the skin(*25*), which could account for the rare skin phenotypes that we experienced. A couple male IKKβKO^Scx^ mice presented with tumor-like growths around their groin area. This may be indicative of a testicular growth, as *Scx* has previous been implicated in Sertoli cell function and differentiation(*26*). These tendon-independent effects are an important reminder that while *Scx* is a highly useful tool for studying tendon biology, healing, and development, *Scx* is expressed by more than just tendon and ligament cells.

One limitation of this study is that the Scx-Cre targets all Scx^Lin^ cells, tendon-derived or not, in a non-inducible manner. Initially this study crossed the IKKβ^F/F^ animals to the inducible Scx-Cre^ERT2^ mouse to generate conditional IKKβ knockout in tendon cells. Unfortunately, insufficient DNA recombination was observed following TAM treatment, and the Scx-Cre mouse was utilized instead. A second limitation is that IKKβ signals through more than just the canonical NF-κB cascade, complicating interpretation of results. Moving forward, the p65^F/F^ mouse(*27*) can be utilized to specifically examine canonical NF-κB signaling in flexor tendon healing.

Another limitation is that no time points beyond day 28 post-repair were examined, making it impossible to know if IKKβKO^Scx^ tendon gliding function would continue to worsen over time or if the excess scar tissue at D28 in IKKβKO^Scx^ healing tendons would eventually remodel successfully into stronger tendon with no negative impact on gliding function. However, the specific human tenolysis sample shown here was obtained 3+years following the initial injury and still exhibited p-p65+ and Bcl2+ myofibroblasts (Figs. 1 and 8), demonstrating the persistence of this finding for several years in the clinical setting.

Taken together, these data suggest that Scx^Lin^ cells activate canonical NF-κB signaling during late stages of tendon healing, where altered canonical NF-κB signaling by Scx^Lin^ cells influences scar-mediated tendon healing and cell survival. Interestingly we also provide evidence that myofibroblasts within human tendon scar tissue have activated canonical NF-κB signaling and express survival signal Bcl-2, suggesting that this finding is clinically relevant. Understanding the cellular contributors to scar-mediated tendon healing and the molecular mechanisms through which they drive scar formation is necessary to design effective therapies to combat excessive scar deposition and drive regenerative healing.

## Materials and Methods

### Study Design

The goal of this study was to investigate the temporal and spatial activation pattern of canonical NF-κB signaling during tendon healing with the goal of delineating cell-type-specific effects of NF-κB knockdown on healing. Genetic mouse models were utilized to determine that tendon cells and myofibroblasts activate and maintain canonical NF-κB signaling throughout healing. We also confirmed that canonical NF-κB-expressing myofibroblasts are present in human tendon scar tissue. We hypothesized that Scx^Lin^ specific knockdown of canonical NF-κB kinase IKKβ would result in decreased scar tissue formation during healing. We investigated these effects via gliding testing, tensile mechanical testing, histology, immunofluorescence, and western blot. Sample size and statistical methods were determined from previous studies and used to inform the experimental design of this work(*13, 17, 18, 28*) in order to determine significant differences in experimental outcomes. Samples for gliding and biomechanical testing, histology and immunofluorescence, and western blotting were randomized, and blinding occurred during analysis when possible. Specifics on statistical methods and sample size are included in the figure legends.

### Animal Ethics

This study was carried out in strict accordance with the recommendations in the Guide for the Care and Use of Laboratory Animals of the National Institutes of Health. All animal procedures described were approved by the University Committee on Animal Research (UCAR) at the University of Rochester Medical Center.

### Mice

Scx-Cre^ERT2^ and Scx-Cre mice were generously provided by Dr. Ronen Schweitzer. IKKβ^F/F^mice(*29*) were generously provided by Dr. Michael Karin. C57Bl/6J (#000664) and ROSA-Ai9^F/F^ (#007909) mice were obtained from the Jackson Laboratory (Bar Harbor, ME, USA). ROSA-Ai9^F/F^ mice express Tomato red fluorescence in the presence of Cre-mediated recombination(*30*). Scx-Cre^ERT2^ and Scx-Cre mice were crossed to the ROSA-Ai9^F/F^ reporter mouse to trace adult Scx+ tendon cells (Scx^Ai9^) and *Scx*-lineage cells (Scx^Lin^), respectively. To label adult Scx+ tendon cells, Scx^Ai9^ animals received three 100mg/kg i.p. tamoxifen (TAM) injections beginning six days prior to tendon injury and repair surgery with a four day washout period between the final injection and surgery, as previously described(*17*). In the IKKβ^F/F^mouse strain, Cre-mediated recombination excises exons 6 and 7 from the *IKK2* locus, resulting in non-functional IKKβ protein(*29*). Scx-Cre mice were crossed to IKKβ^F/F^ animals to generate Scx^Lin^ specific deletion of IKKβ in heterozygous (IKKβHet^Scx^) and homozygous knockouts (IKKβKO^Scx^). All mouse studies were performed with 10-12 week-old male and female mice.

### Human Samples

Collection of human tenolysis tissue samples was approved by the Research Subjects Review Board (RSRB) at the University of Rochester (Protocol #54231).

### Flexor Tendon Repair

Mice underwent complete transection and repair of the flexor digitorum longus (FDL) tendon in the right hindpaw as previously described(*31*). Prior to surgery, mice were injected with 15-20μg of sustained-release buprenorphine. Mice were anesthetized with Ketamine (60mg/kg) and Xylazine (4mg/kg). The FDL tendon was first transected at the myotendinous junction to reduce prevalence of rupture of the repair site and the skin was closed with a 5-0 suture. Immediately afterwards, a small incision was made on the posterior surface of the hindpaw, the FDL tendon was located and completely transected. The tendon was repaired using 8-0 suture and the skin was closed with 5-0 suture. Following surgery, animals resumed prior cage activity, food intake, and water consumption.

### Protein Extraction and Western Blot

Total protein was extracted from wildtype (WT), IKKβHet^Scx^ and IKKβKO^Scx^ littermate uninjured flexor tendons. Protein was also extracted from WT and IKKβKO^Scx^ flexor tendons at days 14 and 28 post-repair. Repaired samples consisted of the repair site and 1-2mm of native tendon on either side of the repair, with three tendons pooled per genotype per timepoint to ensure sufficient protein levels. As such, the protein contribution is reflective of both native tendon tissue and the surrounding scar. Tendons were homogenized using 0.5mm zirconium oxide beads and a Bullet Blender Gold Cell Disrupter (Next Advance Inc., Troy, NY), protein was extracted using radioimmunoprecipitation assay buffer (RIPA) buffer with added protease/phosphatase inhibitors, and 20μg were loaded into each well of a NuPAGE 4-12% Bis-Tris Gel (Invitrogen, Carlsbad, CA). Following transfer, membranes were probed with antibodies for phospho-p65 (1:1000, Cat#: 3033, Cell Signaling, Danvers, MA), total p65 (1:1000, Cat#: 8242, Cell Signaling, Danvers, MA), phospho-ERK1/2 (1:1000, Cat#: 4377, Cell Signaling, Danvers, MA), total ERK1/2 (1:1000, Cat#: 9102, Cell Signaling, Danvers, MA), phospho-JNK (1:1000, Cat#: 4668, Cell Signaling, Danvers, MA), total JNK (1:1000, Cat#: 9252, Cell Signaling, Danvers, MA), phospho-p38 (1:1000, Cat#: 9216, Cell Signaling, Danvers, MA), total p38 (1:1000, Cat#: 9212, Cell Signaling, Danvers, MA), phospho-S6K1 (1:1000, Cat#: 9234, Cell Signaling, Danvers, MA), total S6K1 (1:1000, Cat#: 9202, Cell Signaling, Danvers, MA), phospho-AKT (1:1000, Cat#: 4060, Cell Signaling, Danvers, MA), total AKT (1:1000, Cat#: 9272, Cell Signaling, Danvers, MA), β-catenin (1:1000, Cat#: 610154, BD Biosciences, Franklin Lakes, NJ), p-FOXO3A (1:500, Cat#: ab47285, Abcam, Cambridge, MA) and GAPDH (1:1000, Cat#: 2118, Cell Signaling, Danvers, MA). Full length gels provided in supplemental figures 1 and 2. Band intensity quantification performed using ImageJ and provided in supplemental tables 1 and 2.

### Direct Fluorescence via Frozen Sectioning

Hindpaws from Scx^Lin^ mice were harvested from uninjured contralateral controls and at days 7, 14, and 21 post-repair for frozen sectioning. Hindpaws were fixed in 10% NBF for 24 hours at 4°C, decalcified in 14% EDTA for 4 days at 4°C, and processed in 30% sucrose for 24 hours at 4°C to cryo-protect the tissue. Samples were then embedded in Cryomatrix (ThermoFisher, Waltham, MA, USA) and sectioned into 8μm sagittal sections using a cryotape-transfer method(*32*). Sections were mounted on glass slides using 1% chitosan in 0.25% acetic acid and counterstained with the nuclear stain DAPI. Endogenous fluorescence was imaged on a VS120 Virtual Slide Microscope (Olympus, Waltham, MA, USA). Images are representative of 3-4 specimens per time-point.

### Histology and Immunofluorescence

Hindpaws were harvested at days 14 and 28 post-repair (n = 3-4 per genotype). Hindpaws were fixed in 10% neutral buffered formalin (NBF) at room temperature for 72 hours, decalcified in Webb Jee EDTA (pH 7.2-7.4) for 1 week at room temperature, processed, and embedded in paraffin. All samples were cut into three-micron sagittal sections, followed by de-waxing and dehydrating for analysis. Sections were stained with Alcian blue/hematoxylin and Orange G (ABHOG) for tissue morphology and Masson’s Trichrome for collagen content. For immunofluorescence, sections were probed with antibodies for phospho-p65 (1:200, Cat#: ab86299, Abcam, Cambridge, MA), tdTomato (1:500, Cat#: AB8181, SICGEN, Cantanhede, Portugal), α-SMA-CY3 (1:200, Cat#: C6198, Sigma Life Sciences, St. Louis, MO), α-SMA-FITC (1:500, Cat#: F3777, Sigma Life Sciences, St. Louis, MO), periostin (1:250, Cat#: ab14041, Abcam, Cambridge, MA), F4/80 (1:500, Cat#: sc-26643, Santa Cruz, Dallas, TX), S100a4 (1:2000, Cat#: ab197896, Abcam, Cambridge, MA), Bcl2 (1:500, Cat#: ab182858, Abcam, Cambridge, MA), Bcl-xL (1:400, Cat#: ab32370, Abcam, Cambridge, MA), and Cleaved Caspase 3 (1:100, Cat#: 9661, Cell Signaling, Danvers, MA) overnight at 4°C. The following secondary antibodies were used: donkey anti-rabbit 647 (1:200, Cat#: 711-606-152, Jackson ImmunoResearch, West Grove, PA), Donkey anti-rabbit FITC (1:200, Cat#: 711-546-152), Donkey anti-goat Rhodamine-Red-X (1:200, Cat#: 705-296-147), Donkey anti-rabbit Rhodamine-Red-X (1:200, Cat#: 711-296-152, Jackson ImmunoResearch, West Grove, PA). Sections were counterstained with nuclear DAPI stain and imaged using a VS120 Virtual Slide Microscope (Olympus, Waltham, MA).

### Quantification of Fluorescence

Fluorescent images were processed using Visiopharm image analysis software v.6.7.0.2590 (Visiopharm, Hørsholm, Denmark). Automatic segmentation via a threshold classifier was utilized to define discrete cell populations based on fluorescent channel. Regions of interest were drawn to include the tendon stubs and the entire scar tissue region for processing. The area of each fluorescent signal was calculated, as was the area of each region of interest, and these values were used to determine overall percentage of each cell type. An n=3-4 was used for quantification.

### Gliding Function and Biomechanical Properties Assessment

Tendon gliding function was assessed as previously described(*33*). Hindlimbs were harvested at the knee-joint and the proximal end of the FDL tendon was detached from the myotendinous junction. The FDL tendon was secured between two pieces of tape and was loaded incrementally with small weights ranging from 0 to 19g, with images captured after each load. Measurement of the flexion angle of the metatarsophalangeal (MTP) joint relative to the unloaded position were made using Image J. Gliding resistance was derived from the changes in MTP flexion angle over the range of applied loads. An increase in Gliding Resistance and decrease in MTP Flexion Angle is associated with restricted range of motion and increased scar tissue. Following gliding testing, the FDL tendon was released from the tarsal tunnel. The proximal end of the tendon and the toes of the hindpaw were secured into an Instron 8841 uniaxial testing system (Instron Corporation, Norwood, MA). The tendon was loaded until failure at a rate of 30mm/minute. Eight-14 samples per genotype per time point were analyzed.

### Statistical Analysis

Quantitative data was analyzed via GraphPad Prism and is presented as mean ± standard error of the mean (SEM). Normality was assessed using the Shapiro-Wilk normality test. When data was normally distributed, a student’s t-test was used to analyze data between genotypes at a given time point. When data failed the normality test (gliding function at D28, αSMA at D28, F4/80 at D28), a Mann-Whitney test was used. p values ≤ 0.05 were considered significant.

## Acknowledgements

We would like to thank the Histology, Biochemistry and Molecular Imaging (HBMI) and the Biomechanics, Biomaterials and Multimodal Tissue Imaging (BBMTI) for technical assistance with the histology and biomechanical testing, respectively.

## Funding

This work was supported in part by NIH/ NIAMS K01AR068386 and R01AR073169 (to AEL), F31 AR074815 (to KTB), and R01AR056696 (to HAA). The HBMI and BBMTI Cores were supported by NIH/ NIAMS P30AR069655. The content is solely the responsibility of the authors and does not necessarily represent the official views of the National Institutes of Health.

## Author contributions

Study conception and design: KTB, AEL; Acquisition of data: KTB, EK, CK; Analysis and interpretation of data: KTB, JHJ, HAA, AEL; Drafting of manuscript: KTB, AEL; Revision and approval of manuscript: KTB, EK CK, JHJ, HAA, AEL.

## Competing Interests

The authors declare no competing interests.

## Data Availability Statement

All data generated or analyzed during this study are included in this article.

## Supplemental Material

**Supplemental Table 1. Quantification of uninjured western blot band intensity.**

**Supplemental Table 2. Quantification of repair western blot band intensity.**

**Supplemental Figure 1. Full length western blots for uninjured data.**

**Supplemental Figure 2. Full length western blots for repair data.**

## Supplementary Figures Text

**Supplemental Table 1. Quantification of uninjured western blot band intensity.**

Quantification of western data presented in figure 3. IKKβ band intensity normalized to GAPDH where Uninjured WT band arbitrarily set to 1.

**Supplemental Table 2. Quantification of repair western blot band intensity.** Quantification of western data presented in figure 7. Protein band intensity normalized to GAPDH run on same gel. Band intensity of day 14 WT repair arbitrarily set to 1.

**Supplemental Figure 1. Full length western blots for uninjured data.** Full length western blot data for IKKβ and GAPDH bands presented in figure 3. Quantification of band intensity provided in supplemental table 1.

**Supplemental Figure 2. Full length western blots for repair data.** Full length western blot data for total and p-p65, total and p-ERK1/2, total and p-p38, total and p-JNK, total and p-AKT, total and p-S6K1, β-catenin, and p-Foxo3a presented in figure 7. Quantification of band intensity provided in supplemental table 2.

