## Supplemental Figures for "Tendon Cell Deletion of IKKβ/NF-κB Drives Functionally Deficient Tendon Healing and Altered Cell Survival Signaling In Vivo"

### Slide 1
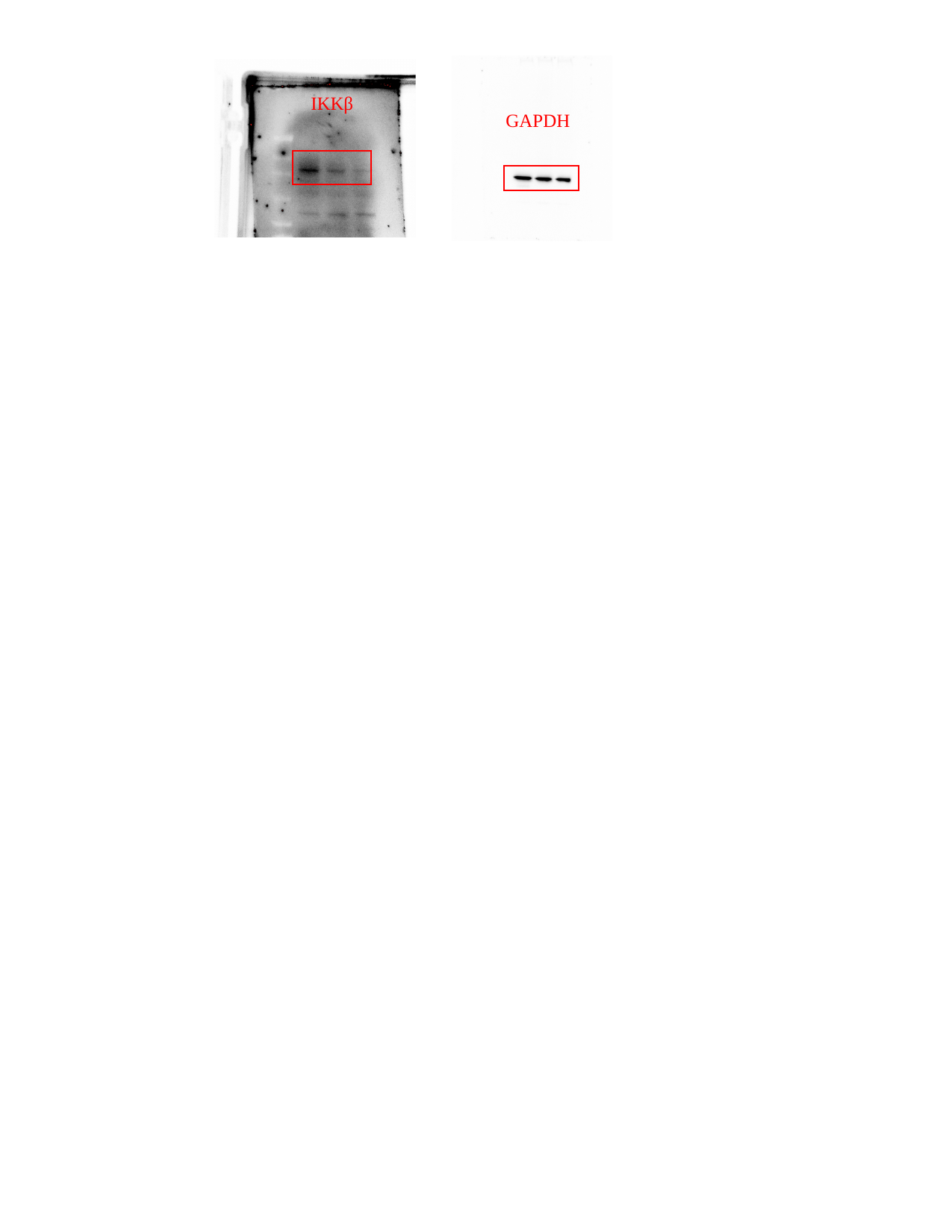

IKKβ
GAPDH

### Slide 2
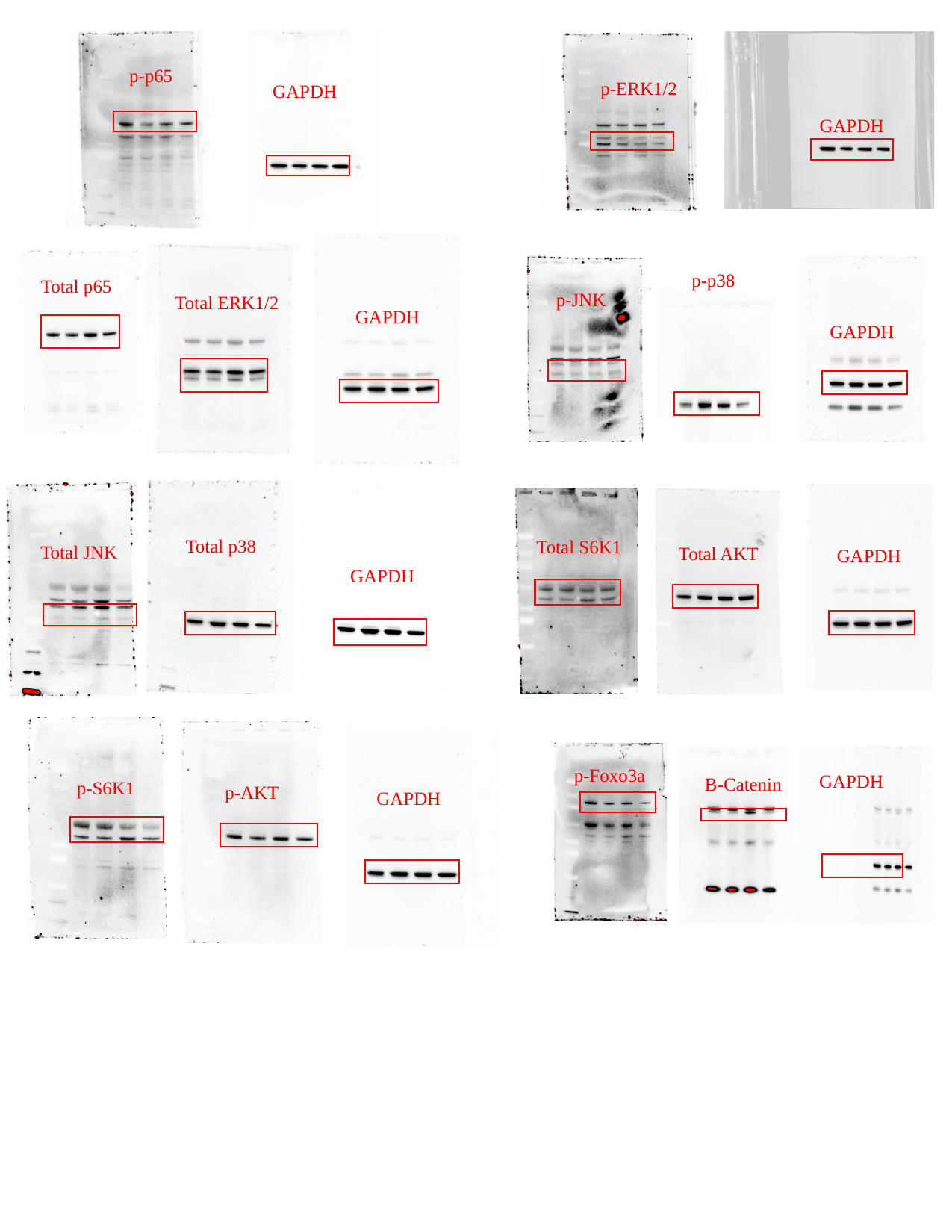

p-p65
p-ERK1/2
GAPDH
GAPDH
p-p38
Total p65
p-JNK
Total ERK1/2
GAPDH
GAPDH
Total p38
Total S6K1
Total JNK
Total AKT
GAPDH
GAPDH
p-Foxo3a
GAPDH
Β-Catenin
p-S6K1
p-AKT
GAPDH

### Slide 3
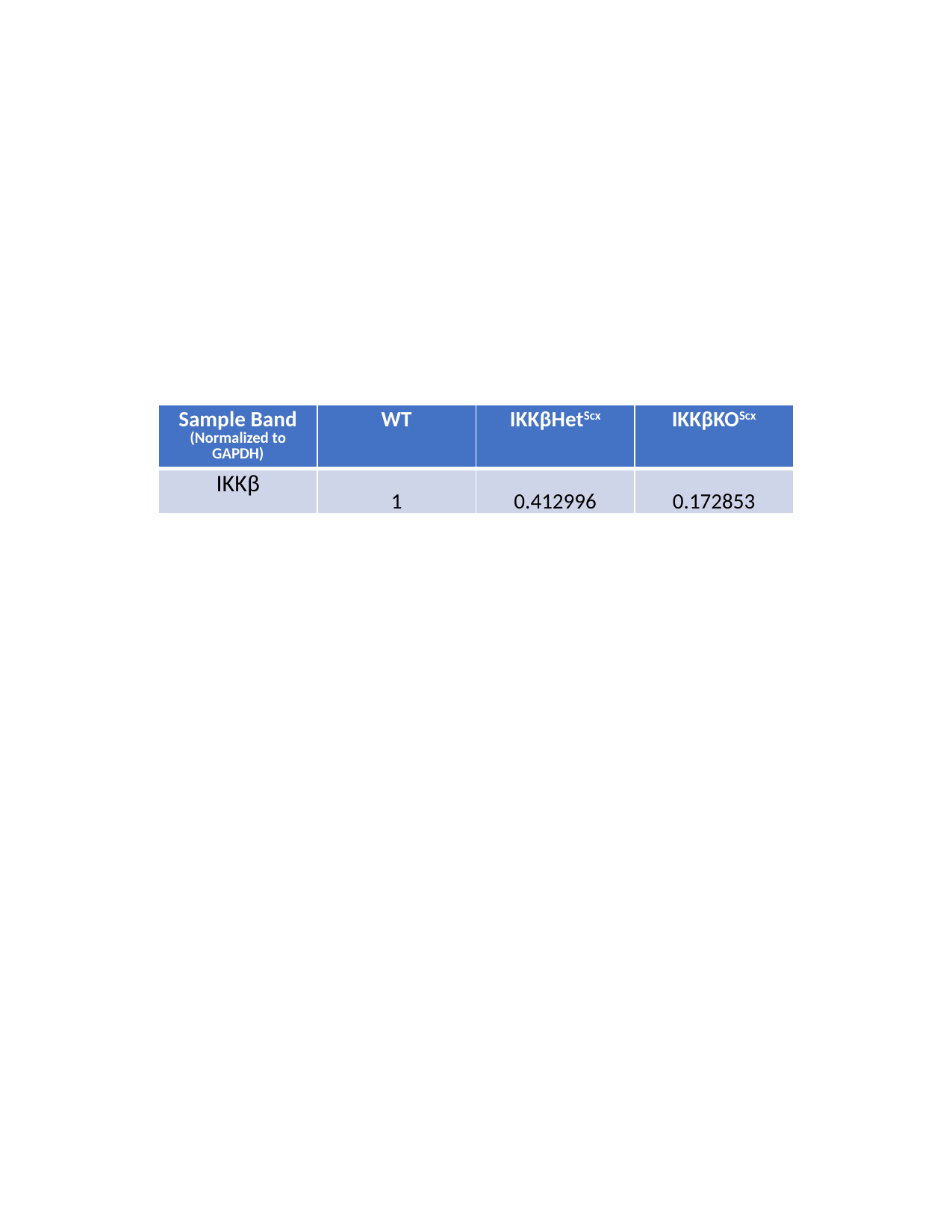

| Sample Band (Normalized to GAPDH) | WT | IKKβHet^Scx^ | IKKβKO^Scx^ |
| --- | --- | --- | --- |
| IKKβ | 1 | 0.412996 | 0.172853 |

### Slide 4
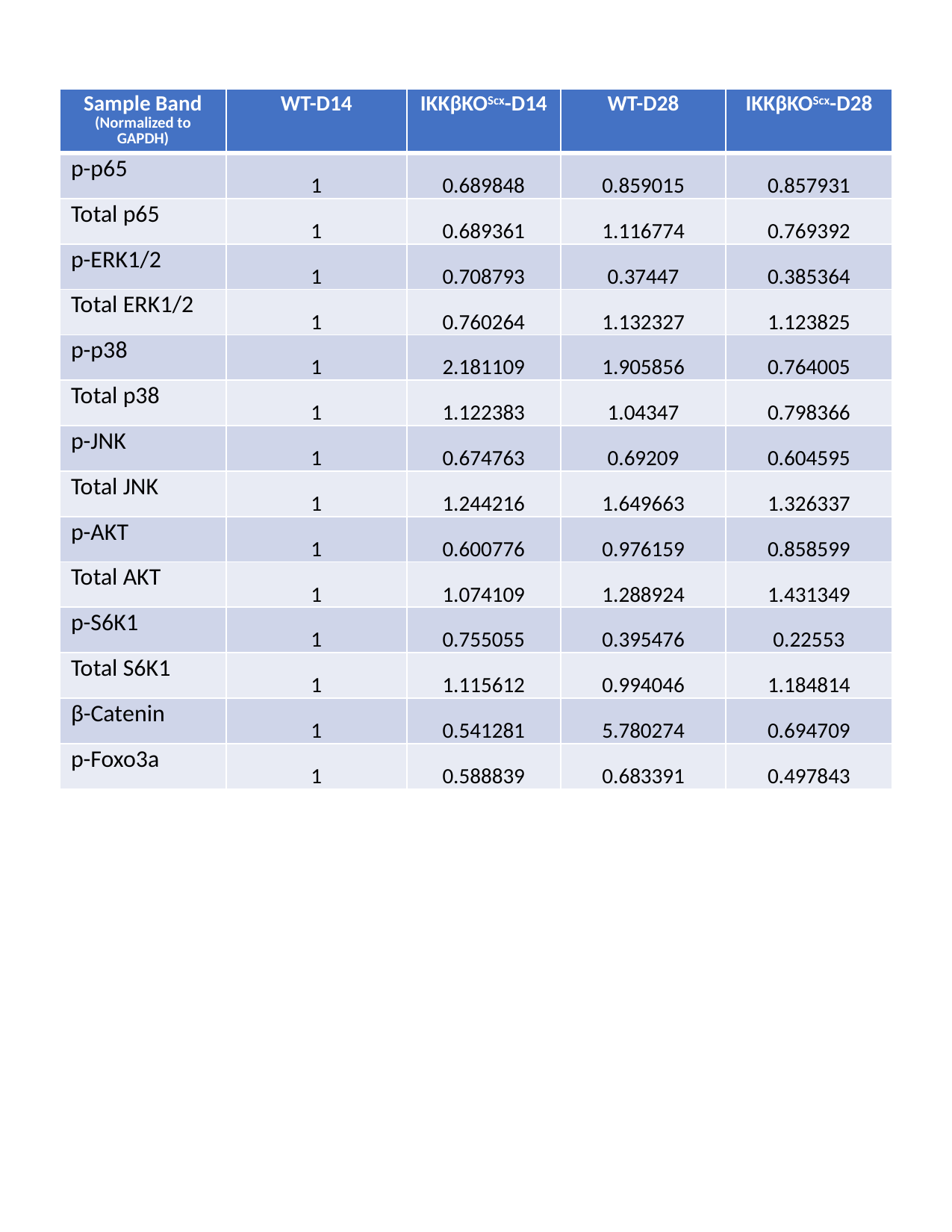

| Sample Band (Normalized to GAPDH) | WT-D14 | IKKβKO^Scx^-D14 | WT-D28 | IKKβKO^Scx^-D28 |
| --- | --- | --- | --- | --- |
| p-p65 | 1 | 0.689848 | 0.859015 | 0.857931 |
| Total p65 | 1 | 0.689361 | 1.116774 | 0.769392 |
| p-ERK1/2 | 1 | 0.708793 | 0.37447 | 0.385364 |
| Total ERK1/2 | 1 | 0.760264 | 1.132327 | 1.123825 |
| p-p38 | 1 | 2.181109 | 1.905856 | 0.764005 |
| Total p38 | 1 | 1.122383 | 1.04347 | 0.798366 |
| p-JNK | 1 | 0.674763 | 0.69209 | 0.604595 |
| Total JNK | 1 | 1.244216 | 1.649663 | 1.326337 |
| p-AKT | 1 | 0.600776 | 0.976159 | 0.858599 |
| Total AKT | 1 | 1.074109 | 1.288924 | 1.431349 |
| p-S6K1 | 1 | 0.755055 | 0.395476 | 0.22553 |
| Total S6K1 | 1 | 1.115612 | 0.994046 | 1.184814 |
| β-Catenin | 1 | 0.541281 | 5.780274 | 0.694709 |
| p-Foxo3a | 1 | 0.588839 | 0.683391 | 0.497843 |
